# Mentoring practices predictive of doctoral student outcomes in a biological sciences cohort

**DOI:** 10.1101/2023.08.18.553806

**Authors:** Reena Debray, Emily A. Dewald-Wang, Katherine K. Ennis

**Author notes:** Corresponding author: Reena Debray.

## Abstract

Despite the importance of a diversity of backgrounds and perspectives in biological research, women, racial and ethnic minorities, and students from non-traditional academic backgrounds remain underrepresented in the composition of university faculty. Through a study on doctoral students at a research-intensive university, we pinpoint advising from faculty as a critical component of graduate student experiences and productivity. Graduate students from minority backgrounds reported lower levels of support from their advisors and research groups. However, working with an advisor from a similar demographic background substantially improved productivity and well-being of these students. Several other aspects of mentoring practices positively predicted student success and belonging, including frequent one-on-one meetings, empathetic and constructive feedback, and relationships with other peer or faculty mentors. Our study highlights the need to renovate graduate education with a focus on retention – not just recruitment – to best prepare students for success in scientific careers.

## Introduction

Despite substantial advances in recent decades, educational attainment outcomes remain uneven across demographic groups in the United States and worldwide (1,2). This is particularly true in STEM fields and presents several problems. First, the composition of people in the scientific workforce influences what types of problems receive attention and how they are approached, which can lead to biases in medical, safety, and other critical research outcomes. For example, male-dominanted engineering teams have historically designed vehicle safety features based on average male proportions, making car crashes more dangerous for female victims than for male victims (3). Second, doctoral training is the primary path to the professoriate, which is tasked with selecting, instructing, and mentoring the next generation of students. If faculty members are most inclined to support students similar to themselves (similarity bias, (4)), or otherwise most effective at supporting students like themselves (5), then barriers to educational access in this generation are likely to repeat in the next.

In conversations about diversifying academic institutions, the vast majority of public attention is given to recruitment (admissions and hiring) (6,7). However, targeting diversity in recruitment is *only* effective and sustainable when also addressing retention (8). In science education, doctoral advising is a major component of the so-called “leaky pipeline” by which students from minority backgrounds leave science. Graduate students from minority backgrounds frequently report less support from their advisors and departments (9–13). These differences in mentoring experiences likely exacerbate existing disparities in the composition of students aspiring to and attaining scientific careers (10,12).

Advisors to doctoral students are tasked simultaneously with many duties: obtaining grant funding to support their students, helping students develop and troubleshoot research projects, helping students network (or networking on their behalf), giving career advice, identifying when students are struggling, and mediating conflicts within research groups. Yet in a 2016 survey, 69% of faculty reported receiving no formal mentorship training (14). Additionally, the power imbalance in academia can lead to a lack of critical feedback from mentees. In the same survey, nearly half of graduate student respondents reported that they had “frequently” experienced poor mentoring, while faculty mentors rarely thought that they mentored poorly (14). Despite these challenges, a growing body of literature indicates that graduate student mentorship is both a skill that can be improved with practice and a science that can be informed by data (15,16).

Effective mentoring can reduce gender- and race-related disparities in graduate student outcomes (10,17). For example, math, physics, and engineering students from minority backgrounds who were enrolled in well-structured and supportive graduate programs were as productive as their white and male peers (12). In this study, we sought to identify specific mentoring practices predictive of graduate student diversity, well-being, and productivity. We administered an online, anonymous questionnaire to doctoral students in four different biological science programs at the University of California, Berkeley. We focused on three major characteristics of the advisor-student relationship: the advisor’s empathy for their student’s needs and concerns, the helpfulness of their feedback, and the frequency and nature of one-on-one meetings. In the case that the student was from a group that has been historically underrepresented in the biological sciences, we asked whether their advisor was also a member of that same demographic group. Lastly, we investigated whether informal mentoring from other faculty members or peers could compensate in the case of inadequate mentoring from the primary dissertation advisor. We collected data on subjective and objective outcomes, including the respondents’ sense of belonging, their self-assessed preparedness for academic and non-academic scientific careers, the number of papers they had published, and their progress towards graduation.

## Results

### Study participants

Participants in this study consisted of graduate students in four biological sciences departments at the University of California, Berkeley: Integrative Biology (IB), Plant and Microbial Biology (PMB), Molecular and Cell Biology (MCB), and Environmental Science, Policy, and Management (ESPM). The study was authorized by the institutional review board at the University of California, Berkeley on March 21, 2022 (protocol number 2022-01-14960). Participants were informed about the goals of the study and plans for data usage and secure storage. They confirmed that they were over 18 years of age, located in the United States, and consented to complete the survey.

Given the low numbers of certain demographic backgrounds in the graduate student population at UC Berkeley, we carefully designed our questions about demographic identity to avoid de-anonymizing participants. This required us to prioritize questions that were the most central to the study goals and to supply broad categories for participants to select from (e.g., “Asian-American and Pacific Islander” rather than specific nationalities). Survey questions were developed in consultation with the UC Berkeley Division of Equity and Inclusion and reviewed by select faculty and graduate student members of the participating departments to ensure the effectiveness and clarity of the survey. A total of 129 students completed the survey, resulting in a response rate of 24% of the membership of the participating departments (**Table S1**). The gender and racial composition of the sample was similar to that of the wider population of participating departments (**Table S2-S3**).

### Areas of need in graduate student mentoring

On average, graduate students reported moderate to high levels of satisfaction with their experiences, with 65-90% of students indicating via an “Agree” or “Strongly Agree” response that their advisors were knowledgeable, helpful, and empathetic, that their dissertation research was meaningful, and that they felt valued within the graduate student community (Fig **S1a**). However, certain aspects of advising consistently fell behind. Just half of graduate students trusted their advisors to take criticism well or felt that conflicts in the lab were handled appropriately. About half of final-year graduate students felt that their program had prepared them adequately for an academic career, but far fewer (28.6%) felt prepared for a non-academic career (Fig **S1b**).

Even when responses were positive on average, the distributions were consistently characterized by a subset of students reporting very low levels of support and inclusion. These students were disproportionately from minority groups – a result that could be the product of direct bias (i.e. microaggressions, macroaggressions, inaccessible events or spaces) and/or indirect effects (students from less privileged backgrounds have often faced other barriers to educational success). Indeed, several of the demographic identities in the survey were correlated. For example, Hispanic or Latino(a/x) students were more likely to identify as multiracial than students of other ethnicities, and students from non-traditional academic backgrounds were more diverse in ages (Fig **1**).

**Figure 1.**
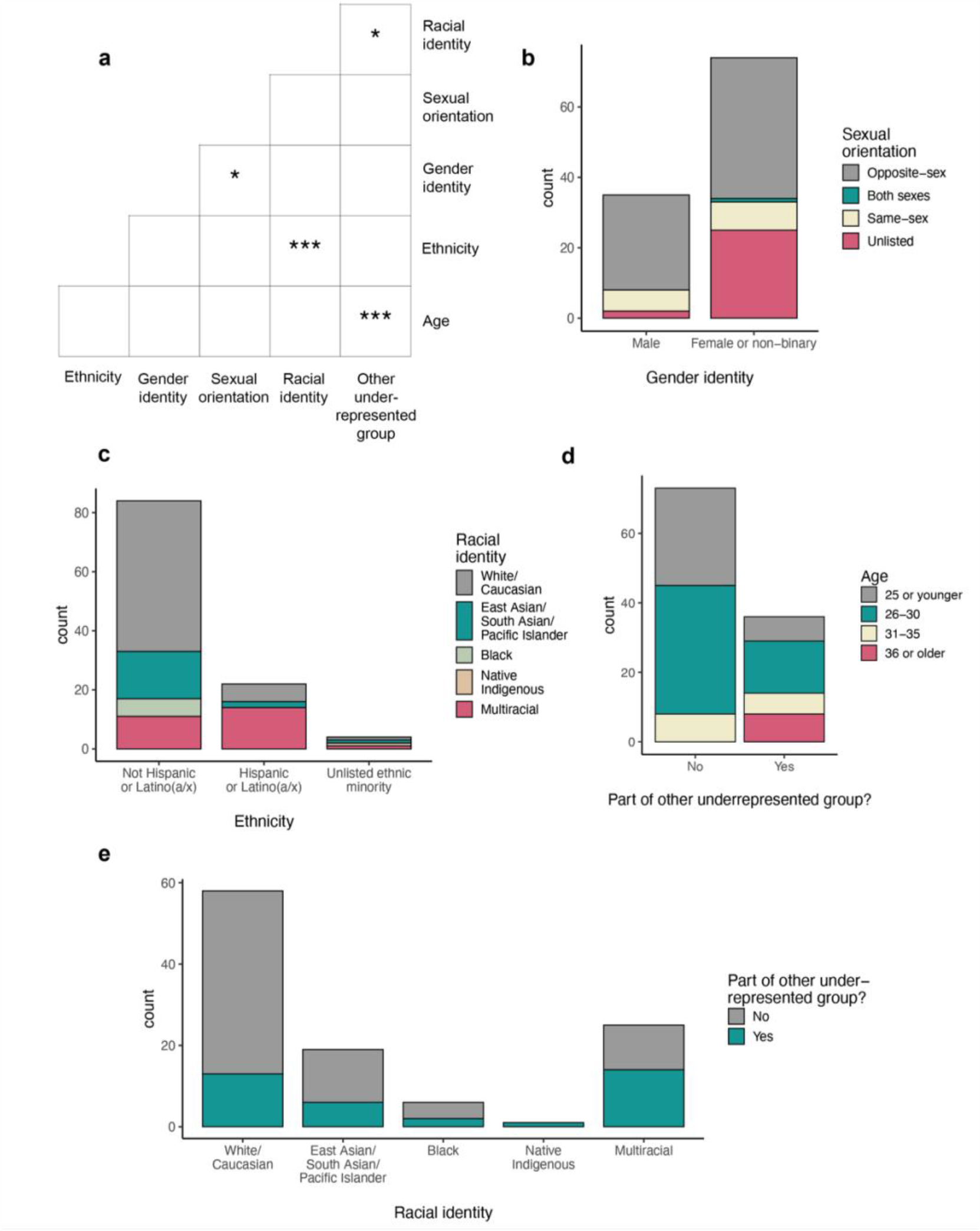
Correlations among demographic groups in this study. **(a)** Demographic variables that were statistically associated according to pairwise chi-squared tests. Asterisks indicate p-values: (***) p ≤ 0.001; (**) 0.001< p ≤0.01; (*) 0.01< p ≤0.05. **(b-e)** Bar plots indicating the composition of demographic groups, excluding non-respondents. “Other underrepresented groups” included parents or caregivers, non-US citizens or permanent residents, first in the family to attend college, disabled students, veterans, neuro-atypical students, and/or students who started their current program more than 5 years after their most recent degree. Categories were grouped during data collection to protect the anonymity of respondents, as each individual category was estimated to apply to a small number of potentially identifiable students.

Compared to their male counterparts, female and non-binary students reported less supportive and inclusive experiences, particularly relating to their research groups (Fig **2a**). Students who started their graduate degree after the age of 30 reported less supportive and inclusive experiences compared to their classmates who started at a younger age, particularly in their relationships with their advisors and research groups (Fig **2b**). Finally, students from non-traditional academic backgrounds, such as parents or caregivers, first-generation students, or students with a disability, reported less support from their advisors than their counterparts who did not identify as such (**Fig S2**). Further, the effects of multiple marginalized identities on student experiences were compounded even beyond their individual effects (Fig **2c-e**). We next sought to identify aspects of mentoring that could remedy these disparities.

**Figure 2.**
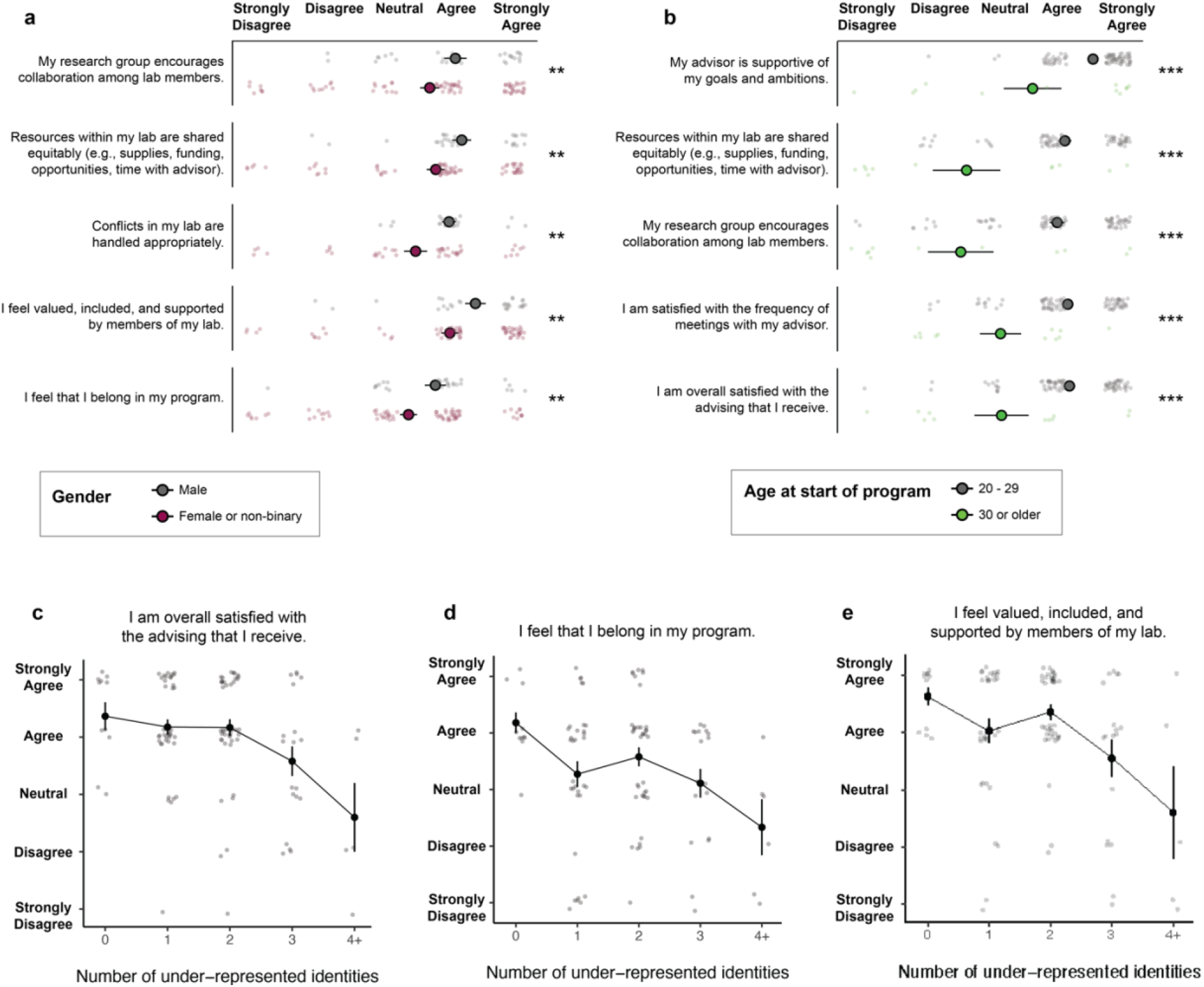
Differences in responses among demographic groups. **(a)** Top five questions with the most disparate responses between male and female/non-binary students. **(b)** Top five questions with the most disparate responses between students who started their graduate degrees before or after the age of 30. Points and error bars represent the mean and standard error, respectively, of Likert responses converted to a 1-5 numerical scale. Statistical significance for each question was measured based on the distribution of differences between members of different groups, which was compared to zero (the null hypothesis, that no differences exist) using a one-sample t-test with degrees of freedom equal to the number of graduate students in the minority category minus one. Asterisks indicate p-values: (***) p ≤ 0.001; (**) 0.001< p ≤0.01; (*) 0.01< p ≤0.05. **(c-e)** Responses to select questions among respondents who identified as multiple underrepresented identities (female or non-binary, started program after age 30, demographic mismatch with advisor, non-traditional academic background, other identity as specified in comments).

### Structure and support

The overall quality of the student-advisor relationship consistently ranked among the top predictors of research progress, self-assessed career preparedness, well-being, and sense of belonging in science in our study. To identify specific aspects of effective advisors, we asked students to evaluate their advisor on a range of qualities, including the extent to which they were supportive (“My advisor is empathetic to my concerns and needs”) or gave constructive feedback (“My meetings are constructive and helpful in setting and achieving my goals”). While most students either rated their advisor highly on both traits (n = 79) or neither (n = 20), enough students fell somewhere in between to evaluate the predictive power of each trait for various graduate student outcomes. Controlling for department and stage of the PhD, students with empathetic advisors felt a stronger sense of inclusion and support in their programs and felt that conflicts were handled more productively within their labs (**Fig S3a**). Students whose advisors gave useful feedback rated their own career preparedness higher and were more likely to feel that their research was meaningful (**Fig S3b**). Interestingly, early- and middle-stage stage students seemed to prioritize empathy in their overall evaluation of their advisors, while late-stage students had a stronger preference for feedback (**Fig S3c**).

We asked students several questions about the scheduling structure, frequency, and helpfulness of their individual meetings with their advisors. Frequent meetings were a positive predictor of nearly every aspect of well-being and productivity measured in the survey (Table **1**). Students who met more frequently with their advisors had a clearer sense of what was expected of them, felt more comfortable bringing concerns to their advisors, and had a stronger sense of belonging in the lab and department. They felt more prepared for post-graduate academic careers and were more likely to be on track to graduate within normative time.

**Table 1.**
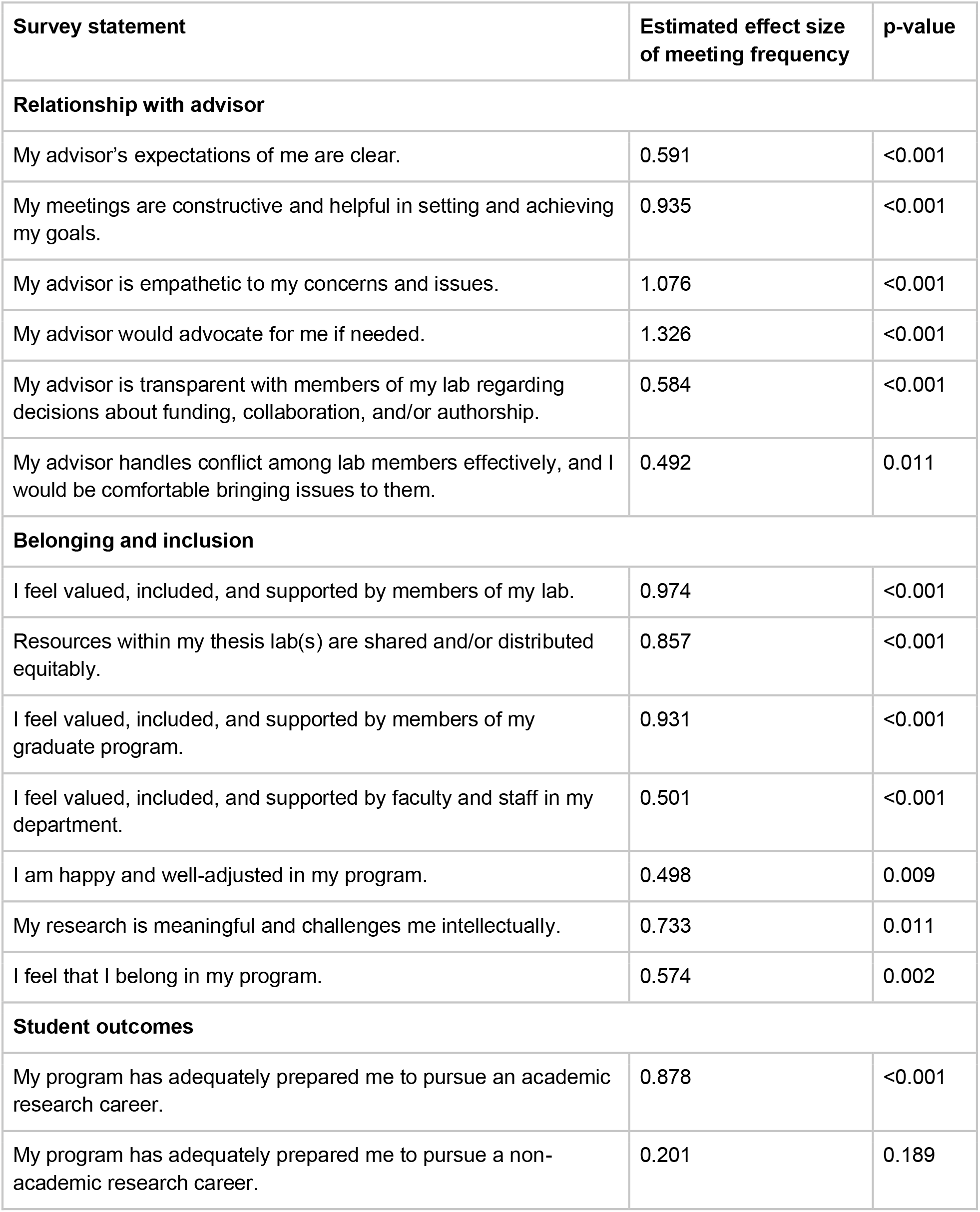
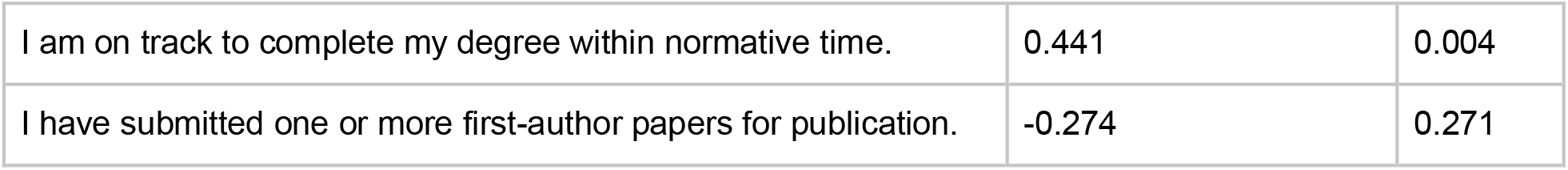
One-on-one meeting frequency positively predicts student experiences and outcomes.

| Survey statement | Estimated effect size of meeting frequency | p-value |
| --- | --- | --- |
| **Relationship with advisor** |  |  |
| My advisor's expectations of me are clear. | 0.591 | <0.001 |
| My meetings are constructive and helpful in setting and achieving my goals. | 0.935 | <0.001 |
| My advisor is empathetic to my concerns and issues. | 1.076 | <0.001 |
| My advisor would advocate for me if needed. | 1.326 | <0.001 |
| My advisor is transparent with members of my lab regarding decisions about funding, collaboration, and/or authorship. | 0.584 | <0.001 |
| My advisor handles conflict among lab members effectively, and I would be comfortable bringing issues to them. | 0.492 | 0.011 |
| **Belonging and inclusion** |  |  |
| I feel valued, included, and supported by members of my lab. | 0.974 | <0.001 |
| Resources within my thesis lab(s) are shared and/or distributed equitably. | 0.857 | <0.001 |
| I feel valued, included, and supported by members of my graduate program. | 0.931 | <0.001 |
| I feel valued, included, and supported by faculty and staff in my department. | 0.501 | <0.001 |
| I am happy and well-adjusted in my program. | 0.498 | 0.009 |
| My research is meaningful and challenges me intellectually. | 0.733 | 0.011 |
| I feel that I belong in my program. | 0.574 | 0.002 |
| **Student outcomes** |  |  |
| My program has adequately prepared me to pursue an academic research career. | 0.878 | <0.001 |
| My program has adequately prepared me to pursue a non-academic research career. | 0.201 | 0.189 |
| I am on track to complete my degree within normative time. | 0.441 | 0.004 |
| I have submitted one or more first-author papers for publication. | -0.274 | 0.271 |

The frequency with which students saw their advisors was closely related to how meeting scheduling was structured. Students whose advisors had an open-door policy or standing appointments ended up meeting more frequently than students whose advisors required them to individually schedule meetings as needed (Fig **3a**). Meeting frequency was also related to the size of the research group, with graduate students in the largest or smallest labs seeing the least of their advisors (Fig **3b**). In large labs, faculty likely face constraints on their time and/or delegate supervision to postdocs and senior graduate students. On the other end of the spectrum, labs headed by faculty who invest less in mentoring might face issues with recruitment and/or retention over time. In support of this second possibility, students who were dissatisfied with their advisors came on average (though not exclusively) from smaller labs (**Fig S4**).

**Figure 3.**
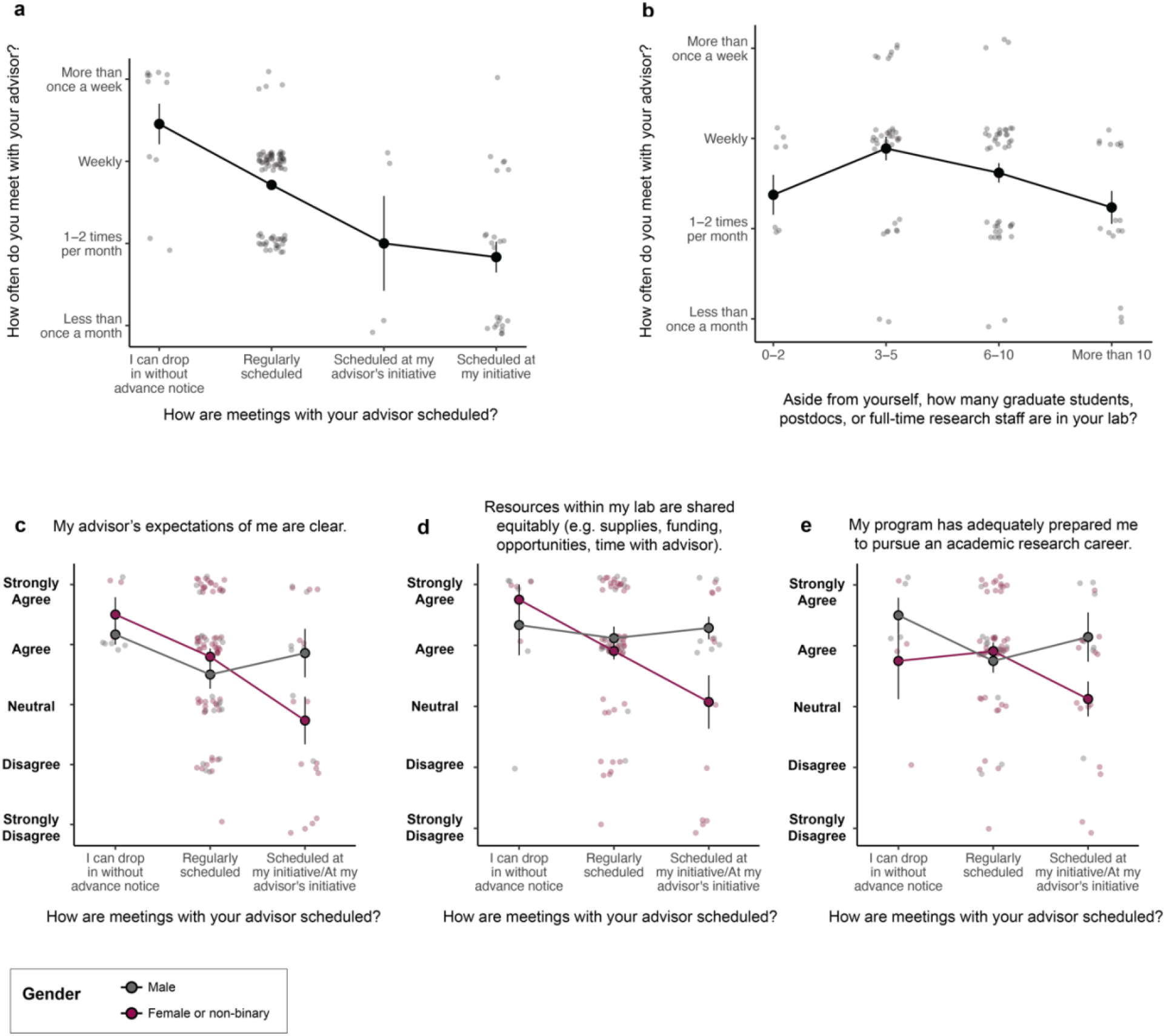
Frequency and scheduling of individual advising meetings. **(a)** Graduate students with as-needed meetings saw their advisors less frequently than students with drop-in or regular meetings (cumulative link mixed mode, p<0.001). **(b)** Graduate students in the largest and smallest labs saw the least of their advisors (nonlinear regression, p=0.007). **(c-e)** Graduate student outcomes according to meeting scheduling format and gender (cumulative link mixed model, p<0.05 for gender x meeting schedule interaction in all cases). Points and error bars represent the mean and standard error, respectively, of Likert responses converted to a 1-5 numerical scale.

Meeting frequency and structure were especially important for graduate students from minority backgrounds. Female and non-binary students who only met with their advisors on an as-needed basis found their advisors less supportive, less transparent about decisions such as funding and authorship, and less clear about expectations compared to female and non-binary students with regular meetings or male students (Fig **3c-e**). Despite publishing at similar rates to their peers, these students were less likely to believe that their dissertation research was meaningful or that they would be prepared for an academic research career after graduate school. Female and non-binary students with as-needed meeting schedules were also the most likely to identify equity issues in their labs, perhaps suggesting that advisors with this system were not equally available to all lab members.

### Representation by advisors

Any student who identified with a demographic that was historically underrepresented in the biological sciences (including race, ethnicity, nationality, gender, sexuality, first-generation status, disability, veteran status, and/or caregiver status) was shown an additional question: *Do you feel that your advisor shares any of those identities?* (Fig **4a**)

**Figure 4.**
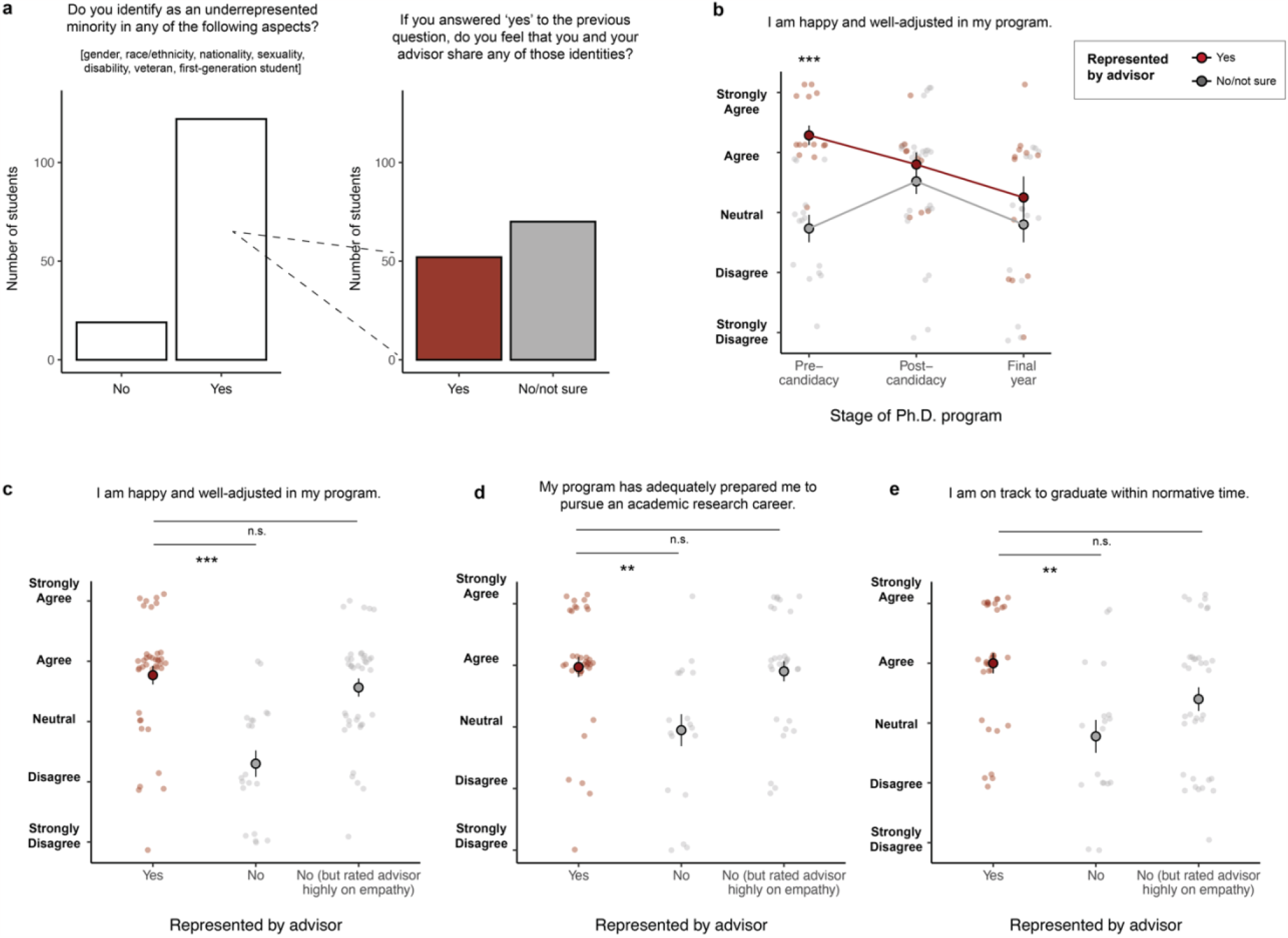
Demographic representation in advisors. **(a)** Proportion of students who identified as an underrepresented minority in any respect, and of those, proportion who worked with advisors of a similar demographic. **(b)** Effect of advisor representation was strongest at the early stages of the PhD. **(c-e)** Graduate student outcomes among students who worked with advisors of a similar demographic, students whose advisors were highly empathetic (answered “Agree” or “Strongly Agree” to the statement “My advisor is empathetic to my concerns and needs”), and students whose advisors were neither. Points and error bars represent the mean and standard error, respectively, of Likert responses converted to a 1-5 numerical scale. Asterisks indicate p-values: (***) p ≤ 0.001; (**) 0.001< p ≤0.01; (*) 0.01< p ≤0.05.

Students who felt represented by their advisors were more likely to feel that their advisors were empathetic and would advocate for them, that they understood what was expected of them, that they belonged in their graduate program, and that their research was meaningful (Table **2**). They felt more prepared for academic and non-academic careers after graduate school. These students were also more likely to have submitted a first-author paper for publication and to be on track to graduate within normative time. The effects of advisor representation on students’ sense of inclusion and belonging were strongest for students early in their programs (Fig **4b**).

**Table 2.** Representation by advisors positively predicts student experiences and outcomes.

| Survey statement | Estimated effect size of meeting frequency | p-value |
| --- | --- | --- |
| **Relationship with advisor** |  |  |
| My advisor's expectations of me are clear. | 1.097 | 0.011 |
| My meetings are constructive and helpful in setting and achieving my goals. | 0.890 | 0.026 |
| My advisor is empathetic to my concerns and issues. | 1.514 | <0.001 |
| My advisor would advocate for me if needed. | 0.954 | 0.035 |
| My advisor is transparent with members of my lab regarding decisions about funding, collaboration, and/or authorship. | 0.190 | 0.621 |
| My advisor handles conflict among lab members effectively, and I would be comfortable bringing issues to them. | -0.354 | 0.397 |
| **Belonging and inclusion** |  |  |
| I feel valued, included, and supported by members of my lab. | 0.785 | 0.056 |
| Resources within my thesis lab(s) are shared and/or distributed equitably. | 0.231 | 0.576 |
| I feel valued, included, and supported by members of my graduate program. | 0.685 | 0.102 |
| I feel valued, included, and supported by faculty and staff in my department. | 0.688 | 0.076 |
| I am happy and well-adjusted in my program. | 1.275 | 0.002 |
| My research is meaningful and challenges me intellectually. | 1.276 | 0.004 |
| I feel that I belong in my program. | 1.210 | 0.002 |
| **Student outcomes** |  |  |
| My program has adequately prepared me to pursue an academic research career. | 1.127 | 0.011 |
| My program has adequately prepared me to pursue a non-academic research career. | 1.019 | 0.013 |
| I am on track to complete my degree within normative time. | 1.462 | <0.001 |
| I have submitted one or more first-author papers for publication. | 1.191 | 0.014 |

Notably, students who did not share any demographic identities with their advisors but rated their advisors highly on empathy tended to be similarly happy and productive as students who did feel represented by their advisors (Fig **4c**). This observation suggests that non-minority advisors can be highly effective mentors to students from minority backgrounds provided that they are open to learning about issues they have not personally experienced.

### Informal mentorship and research community

We asked respondents about the quality and quantity of their connections outside of their formal advising relationship. In general, students reported high levels of support from peers or near-peers on issues pertaining to research practices, scientific careers, and personal issues such as discrimination or work-life balance. Far fewer reported that they had received such advice from faculty, especially on personal issues (**Fig S5a**,**b**).

Many students were affiliated with research organizations within or beyond their departments. For example, graduate students at the University of California, Berkeley whose research focuses on paleontology, entomology, zoology, or botany have the opportunity to affiliate with specialized research museums, affording access to additional seminars, social events, and communal workspaces. Similarly, students working in interdisciplinary research areas such as computational biology have the opportunity to affiliate with graduate groups that hold additional seminars and retreats. Students in either of these types of organizations (but not research institutes) felt more supported and valued in their graduate programs (**Fig S5c**).

Our study identified numerous gaps in student outcomes related to the quality of their relationship with their dissertation advisor. Support and mentorship from other members of the department reduced or closed many of these gaps (Fig **5a-c**). For example, students who felt their advisors would not advocate for them had a lower sense of belonging in science, but this disparity disappeared among those who reported high levels of support from their peers.

**Figure 5.**
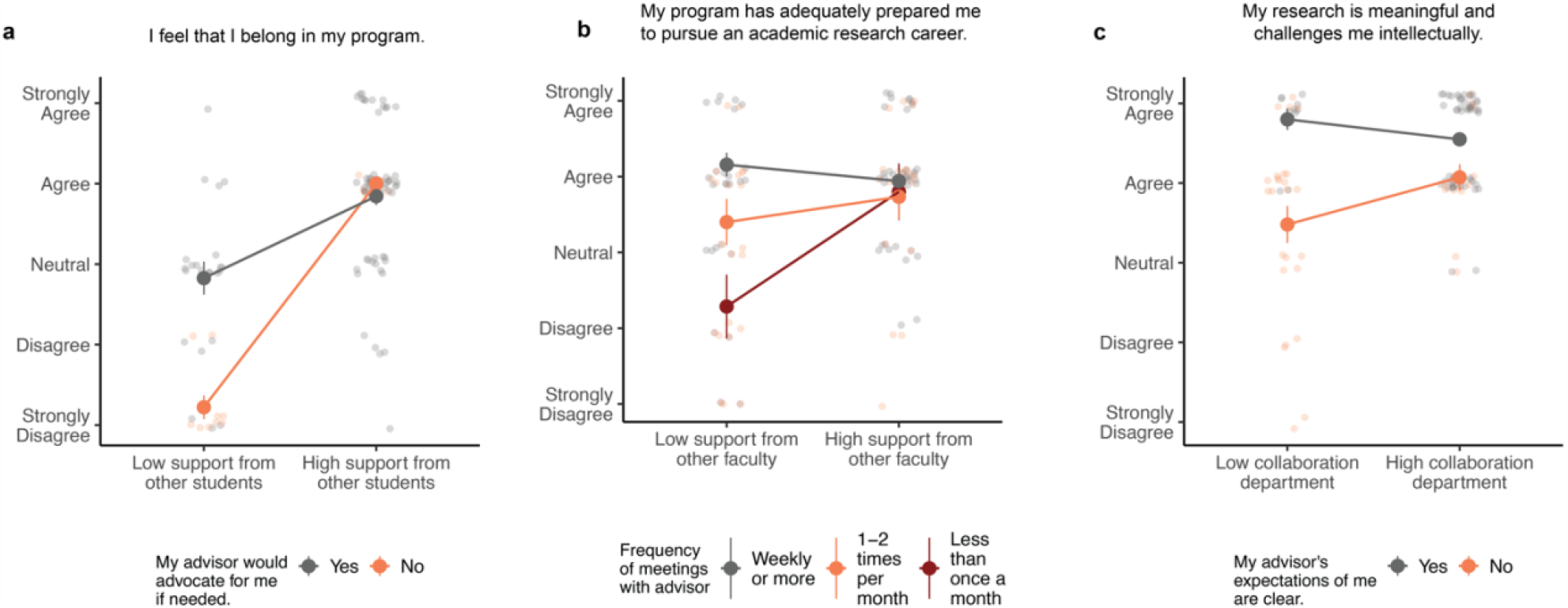
Informal mentorship and community. **(a)** Support from other students appears to compensate for lack of advisor support (cumulative link mixed model, main effect of advisor support p<0.001, main effect of student support p<0.001, interaction effect p=0.008). **(b)** Research advice from other faculty appears to compensate for infrequent meetings with advisor (cumulative link mixed model, main effect of advisor meetings p<0.001, main effect of other faculty p=0.048, interaction effect p=0.078). **(c)** A collaborative departmental culture appears to compensate for less guidance from advisors (cumulative link mixed model, main effect of advisor guidance p<0.001, main effect of collaborative culture p=0.019, interaction effect p=0.048). Points and error bars represent the mean and standard error, respectively, of Likert responses converted to a 1-5 numerical scale. Asterisks indicate p-values: (***) p≤0.001; (**) 0.001< p ≤0.01; (*) 0.01< p ≤0.05.

Students who met infrequently with their advisors felt less prepared for academic careers, but this disparity disappeared among those who reported receiving feedback on their research from other faculty members. Lastly, students whose advisors did not set clear expectations for their progress were less likely to feel that their research was meaningful (perhaps reflecting a perceived lack of investment from their advisor), but being in a department with a collaborative culture helped to narrow this gap.

## Discussion

In this study, we administered an anonymous questionnaire to graduate students in the biological sciences to identify mentoring practices associated with the academic success, diversity, and well-being of respondents. Students from certain groups that have been historically underrepresented in the biological sciences reported less supportive advisors and research groups. Poor advising experiences were not only more common for minority students, but also affected these students’ well-being more negatively. Further, students belonging to multiple minority groups reported even lower levels of support and belonging than those belonging to just one, suggesting additive effects of intersectional identities. We identified individual meetings with advisors, diversity and empathy at the faculty level, and informal mentorship as key interventions to reduce disparities and predict graduate student success and well-being.

### Disparities in student experiences

Consistent with many other reports (9,10,12,18), our study found that female and non-binary graduate students reported lower levels of inclusion, belonging, and equity than their male counterparts. For these students, the support and availability of their advisors was more critical to their confidence and perceived career readiness, even though in reality they published at similar rates to their male peers. Students who came from non-traditional academic backgrounds such as delayed-entry students, first-generation college students, veterans, and parents of young children also reported lower levels of belonging and less support from their advisors. While these disparities are identified in the literature (19,20), they have traditionally attracted less attention than gender from stakeholders or funding agencies, underscoring the importance of developing institutional support systems for unique needs of these groups.

We analyzed participants’ racial identities using either specific categories (e.g., Black; Asian-American) or broader aggregations (e.g., White; Person of Color; Multiracial). In either case, students from underrepresented racial backgrounds did not report substantially different experiences from white students. Why did we not observe an effect of race, despite many previous reports of racial discrimination in scientific workplaces (12,18,21)? There are several possibilities. First, in order to preserve anonymity we deliberately did not collect demographic data at a fine-grained level, though we are aware that designations such as ‘Asian-American’ are not a monolith (21). Second, students who did not respond to the race question at all, or indicated that none of the options we provided captured their identity, were consistently among the unhappiest in the survey, suggesting that some of the most critical populations for this question were not represented because they did not trust the survey (**Fig S6**). Finally, many aspects of racial inequities are indirect and affect certain groups more than others. For example, socioeconomic status and family educational attainment co-vary with race but are not equally relevant to all people of color (17). In our study, students of Black, Native, or multiracial heritage were disproportionately likely to come from non-traditional academic backgrounds, which in turn closely correlated with lower experiences of inclusion and belonging. This observation suggests that identifying race-related disparities in graduate student experiences is best done with survey items that directly target the nature of the opportunity gap in question.

### Structure and support

A common challenge that academic mentors cite is finding the appropriate balance between giving students positive encouragement and critical feedback (22). Yet in our study, empathy and constructive criticism did not trade off; rather, students were always the happiest and most productive when they received both. Steele’s foundational work on stereotype threat (23) found that critical feedback for undergraduate students was “strongly motivating when it was coupled with optimism about their intellectual potential” (24), particularly for Black students, while students receiving unbuffered criticism were discouraged (23). In our study, respondents rated their meetings as more helpful and constructive when they occurred more frequently, possibly suggesting that students view feedback more positively when it comes from someone who regularly invests in them.

Meeting frequency and structure were highly correlated with many other dimensions of success and belonging. This was true for all students, but women and non-binary students in particular were more anxious about navigating their advisors’ expectations in less structured lab environments. This is consistent with previous research that women are less socialized than men to ask for opportunities and resources in the workplace (25,26). These students were also the most likely to identify equity issues within their labs. This perhaps suggests that without an explicit structure in place, advisors invested less in them than in their male colleagues, or that students from less privileged backgrounds may be conditioned to expect that the bar is held higher for them. Our results suggest that frequent, regularly scheduled individual meetings and clear expectations from advisors set a baseline for equitable treatment and help students feel supported in navigating both research-related and personal challenges.

### Representation by advisors

In our study, as observed previously (10,27–29), students from underrepresented demographic backgrounds had a stronger sense of belonging and were more productive (including in their number of publications) when mentored by an advisor who was from a similar background. This effect in our study was strongest for early-stage students, who may not have had many other role models yet.

Multiple mechanisms could explain these findings, all mutually compatible. Students who have more in common with their advisors may feel more confident that people ‘like them’ can succeed (10). They may find more effective solutions to personal difficulties, such as balancing work with raising a family or navigating financial stress, with the help of mentors who have been through similar situations in the past. Finally, it is possible that the operative variable here is not representation *per se*, but that faculty from these backgrounds tend to be more conscientious mentors to all of their students because of the barriers they have faced themselves. To disentangle effects of representation from faculty experience in our dataset would require knowledge of which non-minority students have advisors from minority backgrounds – a level of information that would risk de-anonymizing respondents. However, it is worth noting that in a previous study of graduate student and advisor pairs in the biological sciences, both male and female students were more productive when working with female advisors than with male advisors (29).

Studies such as Pezzoni et al. 2016 have been cited to advocate for increasing diversity at higher levels of education, yet until that is fully realized, faculty members from these backgrounds will continue to be disproportionately tasked with mentorship requests (24). How can we improve mentorship for students from minority backgrounds without over-exerting the small number of minority faculty in each department? Our study found that minority students who did not work with an advisor of the same demographic, but nevertheless rated their advisor highly on empathy, felt as supported and productive as students whose advisors were a closer demographic match. Beyond simply ‘empathy’, this may encompass allocating time and funding equitably within their labs, tailoring their mentoring strategy to each student’s needs, modeling ethical behavior, welcoming honest feedback without fear of repercussions, and advocating for institutional change that supports the needs of underrepresented students (13,14,16,30,31).

### Informal mentorship and research community

In the biological sciences, financial support and recommendation letters are typically provided primarily by the dissertation advisor. However, other mentoring duties such as advice on research, assistance with networking, and support with personal challenges may be filled by the advisor and/or by other mentors or peers. Accordingly, institutional support has been linked to publication record in graduate school, persistence within STEM fields, and job placement after graduate school (12,17,32). Further, students who are dissatisfied with their primary advisors are more likely to seek out secondary mentors (18). Building on these observations, we hypothesized that informal mentors would be the most important for students with low levels of support from their dissertation advisors. Indeed, support and mentorship from the surrounding research community consistently reduced or closed disparities for students who had less helpful experiences with their dissertation advisors. Our dataset further showed that students who belonged to small research organizations within departments and/or interdisciplinary research groups across departments had a stronger sense of community than unaffiliated students, suggesting potential avenues for departmental restructuring that encourage informal mentoring networks.

### Caveats and conclusions

It is important to note the limitations of conclusions that can be drawn from this type of study. These data are both cross-sectional and observational, meaning that any causal conclusions necessarily rely on assumptions about directionality. For example, in this study, graduate students who felt that they belonged in their program were more likely to have completed a first-author paper. But does a sense of scientific identity make students more engaged with their research, or do students who publish subsequently become more confident that they belong? Repeated, identifiable data from the same students over time would be required to find out. With this in mind, we have focused the paper primarily on tangible aspects of the mentoring experience, such as meeting frequency and the demographic backgrounds of advisors, that are less susceptible to reverse causality.

Second, the sample draws exclusively from four biological sciences departments at a single research-intensive university in California. The second point is mitigated in part by the fact that the students themselves are from geographically diverse backgrounds; at the time this survey was administered, 72% of new doctoral students at the university came from outside of California (33). However, to what extent the conclusions presented here apply to graduate students in other fields or at doctoral universities with lower research activity is less clear. Graduate students in the biological sciences typically work in labs that are funded by their advisors’ grants, while in the social sciences and humanities, advising relationships are more characterized by individual meetings that focus on the student’s independent work. Accordingly, doctoral students in the biological and physical sciences reported that their advisors provided more instrumental support than students in the social sciences and humanities, but less intellectual or emotional support, less availability, and less respect (18). Graduate student experiences vary widely by institution types, as well. Students at “R1” universities such as Berkeley receive disproportionate resources such as national fellowships (34), but often have a less diverse student body and may foster less supportive environments. In chemistry, for example, the desire of female graduate students to finish their degree and remain in science declined sharply with departmental prestige (with no such dropoff for men) (10).

Finally, in some instances we relied on self-reported data to assess graduate student outcomes. However, where we were able to benchmark self-assessments against quantitative measures of progress (i.e. publication record), the two were highly correlated (**Fig S7**). This suggests that students are generally well aware of typical progress for graduate students at their career stage.

Katz and Harnett wrote that “Graduate student relations with members of the faculty is regarded by most graduate students as the single most important aspect of the quality of their graduate experience; unfortunately, many also report that it is the single most disappointing aspect of their graduate experience” (35). Our dataset highlights this immense role that doctoral advisors play in the productivity and well-being of their students. Even at a relatively diversity-emphasizing and well-funded institution such as the University of California, Berkeley, students from underrepresented demographic groups still face disproportionate barriers to their success. Our study identifies availability of advisors to their students, diversity at the faculty level, and the support of informal and peer mentors as key attributes that support graduate students and help to remedy these disparities.

## Materials and Methods

### Survey instrument

The survey consisted of 10 questions about the student’s background, including which program they were enrolled in, their approximate stage in the program, and their demographic identity, 37 questions about interactions with their advisor, research group, and department, and 30 questions about their research progress, productivity, and subjective well-being. The majority of questions about the student’s experience with their advisor, lab, and department were presented as a five-point Likert scale (1 = Strongly Disagree, 2 = Disagree, 3 = Neutral, 4 = Agree, 5 = Strongly Agree). Survey items were coded such that a higher score indicated a more positive experience within the academic community (with the exception of the two statements “Travel or lab density restrictions imposed by COVID-19 reduced my research productivity during the past two years” and “Personal or financial challenges imposed by COVID-19 reduced my research productivity during the past two years”). The full survey instrument is available in the Supporting Information (**Table S4**).

### Survey administration

The survey was administered confidentially using the Qualtrics platform. It was distributed electronically via email to all graduate students in the participating departments and remained open from April 26 to May 31, 2022. Completion of the survey was entirely voluntary. Survey participants were encouraged to fill out a separate, unlinked form with their contact information for entry into a drawing for a gift card.

### Statistical analysis

The following data cleaning and formatting steps were taken prior to analysis: 1) Due to the low number of non-binary students in the sample, they were analyzed together with the female students unless they had specifically indicated “Male/Non-binary” as their gender identity. 2) The stage of the respondent’s PhD was defined as either early (has not yet advanced to candidacy, which typically takes place in year 2 of all participating programs), middle (advanced to candidacy, but not graduating within one year), and late (graduating within one year), 3) The approximate age at which the respondent had started their PhD was estimated based on their reported age group and stage of the program. Anyone who was younger than 30 at the time of the survey must have started their PhD before the age of 30. Respondents who were 31-35 and in the pre-candidacy stage were estimated to have started at age 30 or later, and respondents at any stage of their PhD who were 36 or older were estimated to have started at age 30 or later.

Questions about students’ experiences and outcomes were either recorded as Likert scale data (Strongly Disagree, Disagree, Neutral, Agree, Strongly Agree) or another ordinal variable (e.g., whether respondents had published a first-author paper “Never”, “Once”, or “More Than Once”). Parametric tests are not ideal for such data because they assume that the outcome variables were continuous with even spacing between levels (36). Instead, cumulative link mixed models were fitted using the function *clmm* in the R package ordinal (v.2022.11-16) (37). Unless otherwise specified, models included department and stage of PhD as random effects and assumed a logistic distribution. Likert scale data was assumed to be symmetric (that is, the distances between “Strongly Agree” and “Agree” or “Agree” and “Neutral” are not known, but “Strongly Agree” and “Strongly Disagree” are equally far from “Neutral”), while all other ordinal data was fitted with a flexible parameter that did not make any assumptions about symmetry or spacing.

## Data availability

The data collected in this study is potentially identifying and can only be shared upon reasonable request. Researchers interested in investigating or replicating the analyses in this publication can contact the Office for the Protection of Human Subjects at the University of California, Berkeley: 1608 Fourth Street, Suite 220, Berkeley CA, 94710. Phone: (510) 642-7461.

## Acknowledgments

The authors would like to thank the members of the DEI Small Grants Group (Juan Liu, Cindy Looy, Chris Martin, Onja Razafindratsima, Peter Sudmant, Rebecca Tarvin, Jack Tseng, Monica Albe) for providing seed funding from the Life Sciences Faculty Diversity Initiative and feedback on the proposed research. Rebecca Tarvin provided additional support on securing and adhering to the Institutional Review Board protocol. Andrew Eppig provided support on developing and administering the survey instrument. The graduate student, postdoctoral, and faculty members of the participating departments provided additional feedback on interpreting and presenting the results.

