## Supplemental Information for "Mentoring practices predictive of doctoral student outcomes in a biological sciences cohort"

### Supplemental Material for: Mentoring practices predictive of doctoral student outcomes in a biological sciences cohort

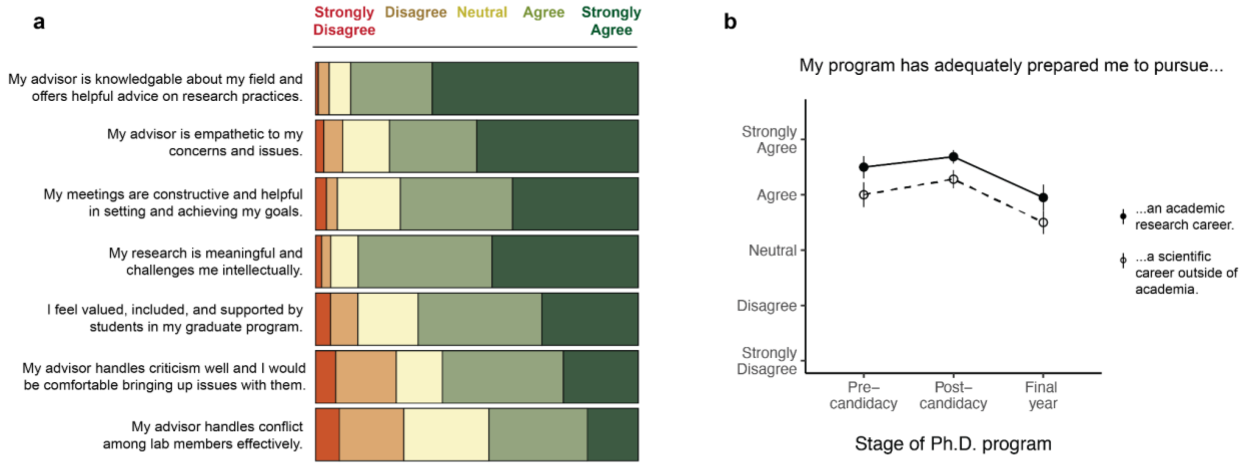

**Figure S1. Overview of graduate student responses to mentorship survey. (a)** Categorical breakdown of responses to select survey items. **(b)** Self-assessed preparedness for academic and non-academic careers. Points and error bars represent the mean and standard error, respectively, of Likert responses converted to a 1-5 scale.

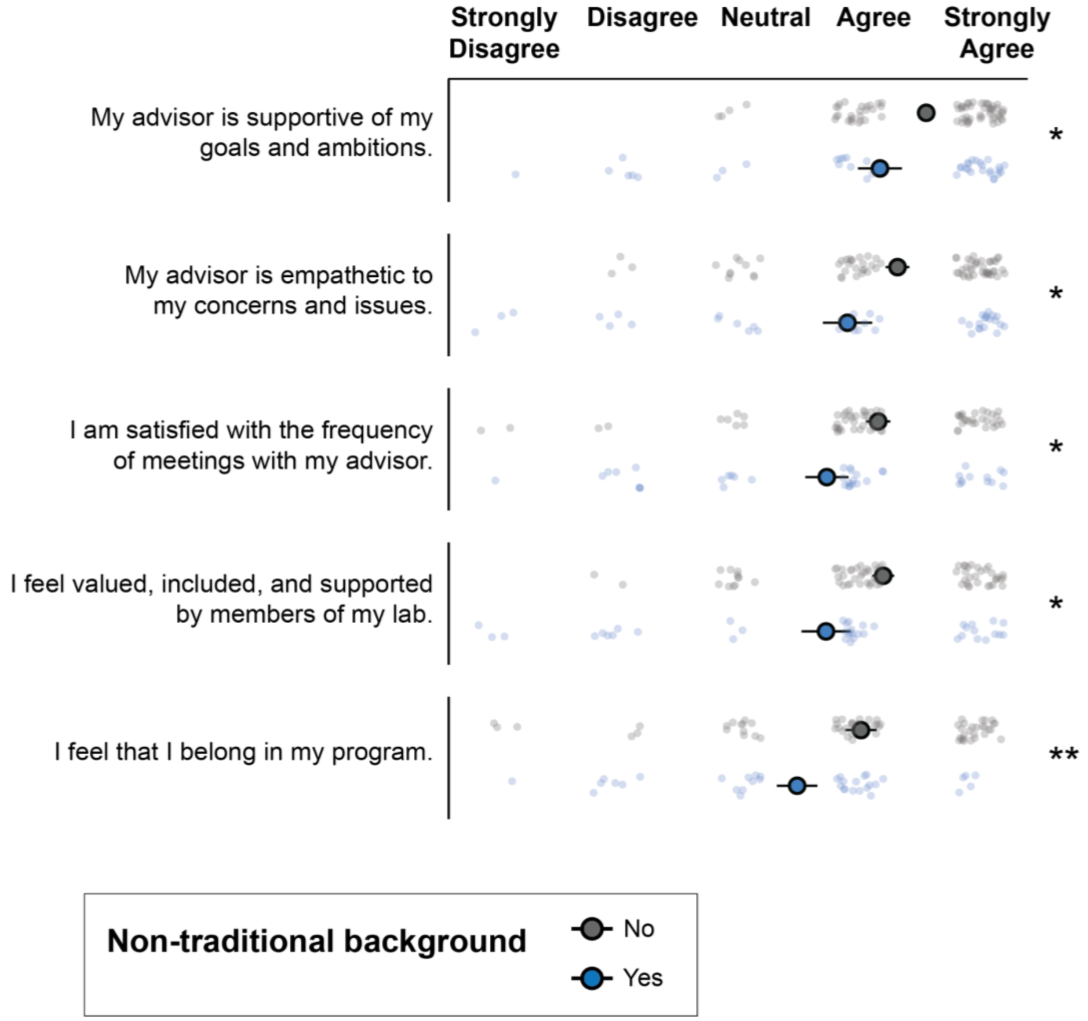

**Figure S2. Differences in responses among demographic groups.** Top five questions with the most disparate responses between students from traditional and non-traditional backgrounds (parent or caregiver, non-US citizen or permanent resident, first in family to attend college, disabled, veteran, neuro-atypical, started program more than 5 years after the most recent degree). Categories were grouped during data collection to protect the anonymity of respondents, as each individual category was estimated to apply to a small number of potentially identifiable students. Points and error bars represent the mean and standard error, respectively, of Likert responses converted to a 1-5 scale. Statistical significance for each question was measured based on the distribution of differences between members of different groups, which was compared to zero (the null hypothesis, that no differences exist) using a one-sample t-test with degrees of freedom equal to the number of graduate students in the minority category minus one. Asterisks indicate p-values: (***) $p \leq 0.001$; (**) $0.001 < p \leq 0.01$; (*) $0.01 < p \leq 0.05$.

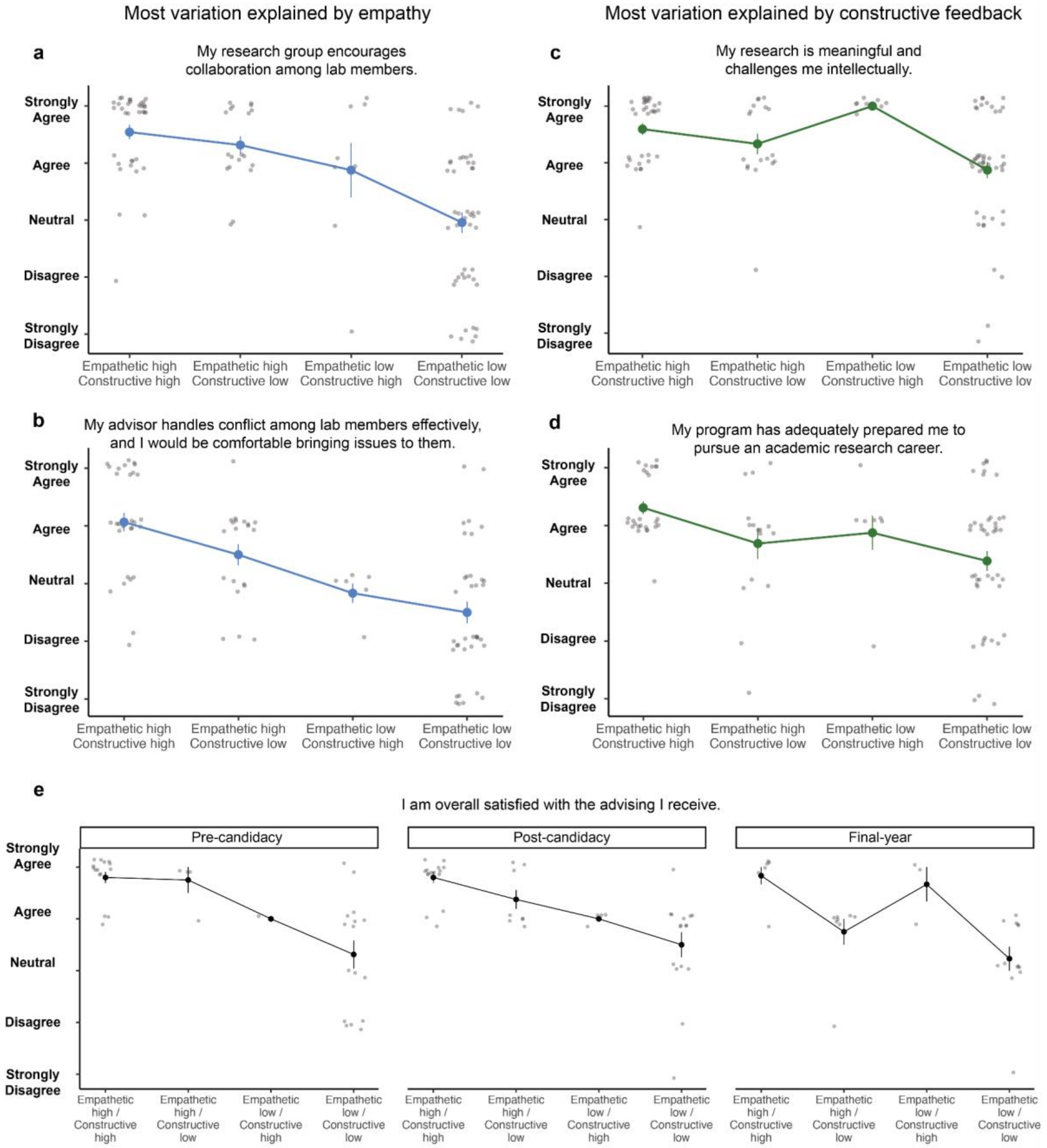

**Figure S3. Predictive value of advisor empathy and constructive feedback. (a-d)** Graduate student outcomes according to whether their advisors scored above or below the median rating on empathy and constructive feedback. **(e)** Overall satisfaction with advising as a function of empathy and constructive feedback at different stages of the PhD. Points and error bars represent the mean and standard error, respectively, of Likert responses converted to a numerical scale (1 = Strongly Disagree, 2 = Disagree, 3 = Neutral, 4 = Agree, 5 = Strongly Agree). Colors indicate which variable was a stronger predictor in a cumulative link mixed model controlling for department and stage of PhD.

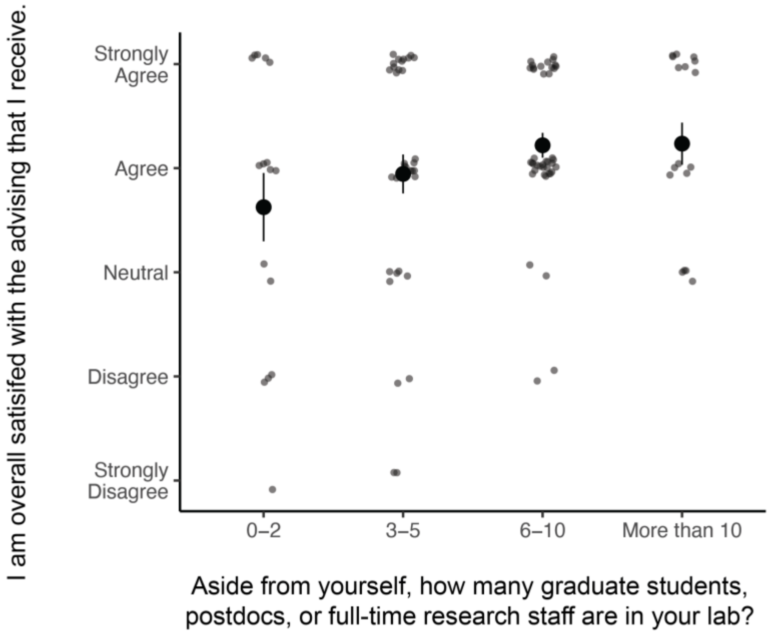

**Figure S4. Relationship between lab size and satisfaction with advising.** On average, students who were dissatisfied with their advisors came from smaller labs. Points and error bars represent the mean and standard error, respectively, of Likert responses converted to a 1-5 scale.

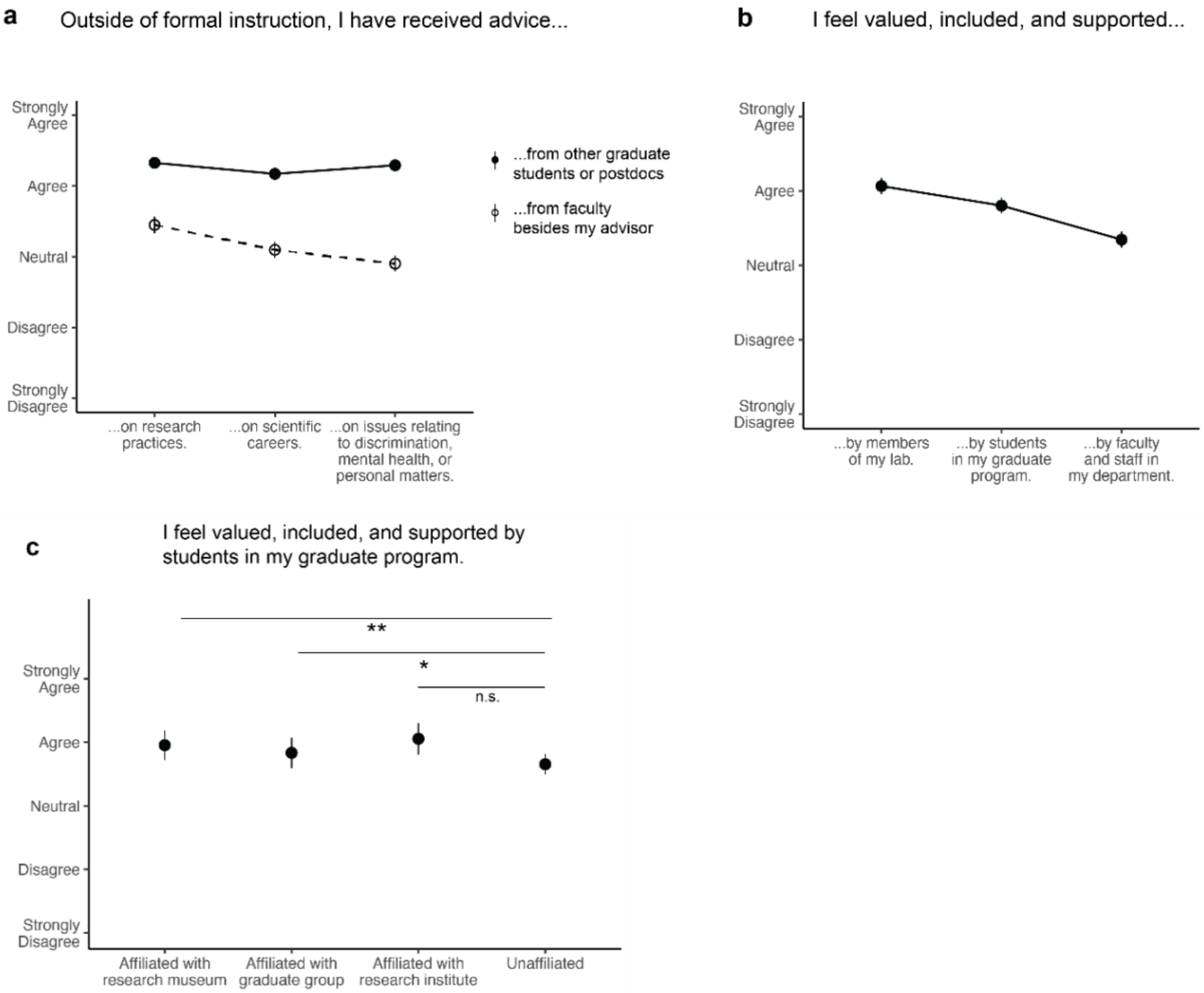

**Figure S5. Sense of belonging and community in the biological sciences. (a)** Research, career, and personal advice from students and faculty. (**b)** Sense of inclusion within and outside the lab. **(c)** Sense of inclusion for students according to lab affiliation. Points and error bars represent the mean and standard error, respectively, of Likert responses converted to a 1-5 scale.

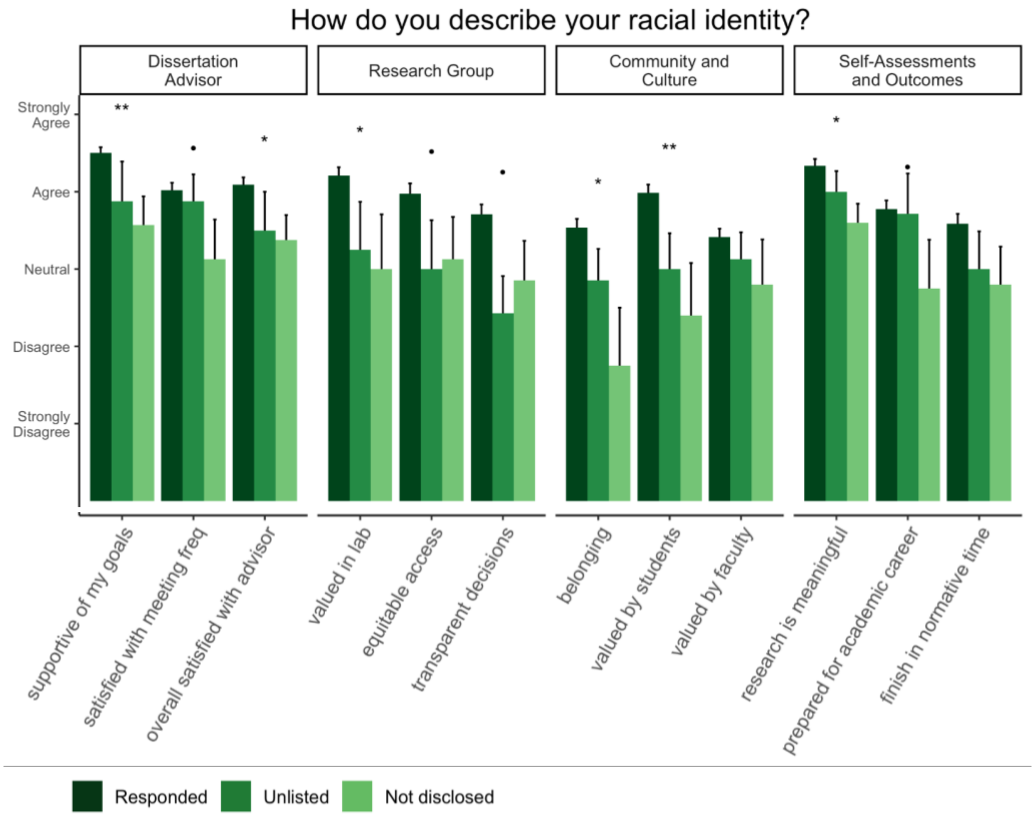

**Figure S6. Experiences of non-respondents to demographic questions.** Responses to various advising experience and well-being questions of students who indicated a specific category on the racial identity (“Responded”), indicated that their identity was not captured by the options provided (“Unlisted”) or elected not to answer the question (“Prefer not to disclose”). Bars and error bars represent the mean and standard error, respectively, of Likert responses converted to a 1-5 scale.

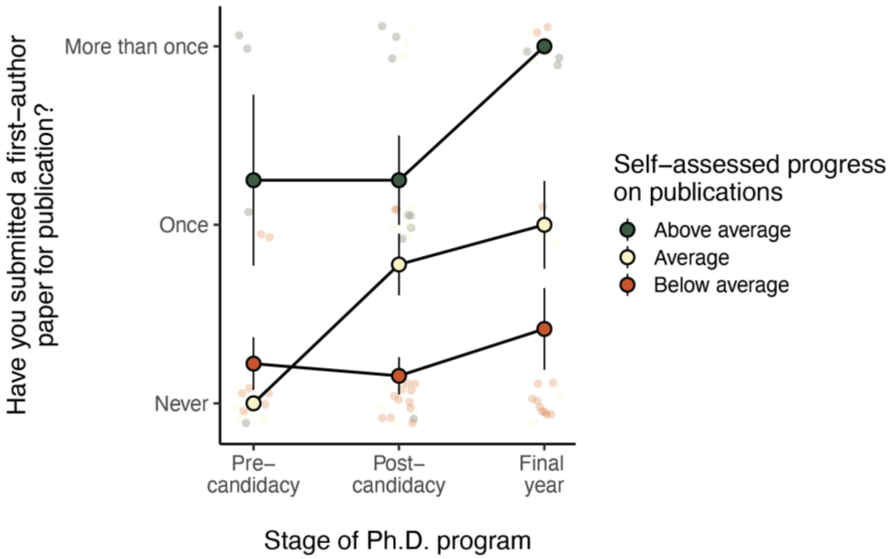

**Figure S7. Correspondence between self-assessed and actual publication rates.** Self-assessed progress on publications was closely related to actual measured progress when accounting for program and stage of Ph.D.

**Table S1. Sample sizes and response rates by department. Enrollment is based on Spring 2022 data from the UC Berkeley Office of Planning and Analysis.**

| Department | Our study | UCB enrollment | Estimated response rate |
|---|---|---|---|
| Integrative Biology (IB) | 42 | 114 | 36.8% |
| Molecular and Cell Biology (MCB) | 41 | 215 | 19.1% |
| Plant and Microbial Biology (PMB) | 24 | 92 | 26.1% |
| Environmental Science, Policy, and Management (ESPM) | 22 | 117 | 18.8% |

**Table S2. Sample sizes and response rates by gender. Enrollment is based on Spring 2022 data from the UC Berkeley Office of Planning and Analysis.**

| Gender identity (this study) | Composition of this study | Gender identity (UCB categories) | Composition of participating departments |
|---|---|---|---|
| Female | 56.6% | Female | 58.0% |
| Male | 27.1% | Male | 41.6% |
| Non-binary<br>Female, Non-binary<br>Male, Non-binary | 5.4%<br>3.1%<br>0.8% | Non-binary | 0.4% |
| NA / prefer not to disclose | 5.4% | | |

**Table S3. Sample sizes and response rates by race. Enrollment is based on Spring 2022 data from the UC Berkeley Office of Planning and Analysis.**

| Racial identity (this study) | Composition of this study | Racial identity (UCB categories) | Composition of participating departments |
|---|---|---|---|
| White/Caucasian | 50.4% | White/Other | 54.0% |
| East Asian/South Asian/Pacific Islander | 16.3% | Asian | 16.3% |
| Black<br>Native/Indigenous | 4.7%<br>0.8% | International<br>Underrepresented Minority | 10.4%<br>20.0% |

| | |
|---|---|
| Multiracial | 10.1% |
| NA / prefer not to disclose | 9.5% |

**Table S4. Full survey instrument disseminated to participating departments.**

| **I. DEMOGRAPHICS** | | | |
|---|---|---|---|
| **Question number** | **Question type** | **Text of question/ statement** | **Options provided to respondents** |
| 1 | Select one | Which of the following graduate programs are you enrolled in? | Integrative Biology; Plant and Microbial Biology (Plant Biology or Microbiology); Molecular and Cell Biology; Environmental Science, Policy, and Management; I am not enrolled in any of these programs. |
| 2 | Select one | Have you advanced to candidacy? | Yes; No; Not applicable (I have graduated or am no longer enrolled as a student) |
| 3 | Select one | Are you planning to graduate within the next 12 months? | Yes; No |
| 4 | Select all that apply | How do you describe your gender identity? | Man; Woman; Unlisted gender minority; Prefer not to disclose |
| 5 | Select all that apply | How do you describe your sexual orientation? | Heterosexual; Homosexual; Unlisted sexual minority; Prefer not to disclose |
| 6 | Select all that apply | How do you describe your racial identity? | White/Caucasian; Black; East Asian/South Asian/Pacific Islander; Native/Indigenous; Unlisted racial minority; Prefer not to disclose |
| 7 | Select all that apply | How do you describe your ethnicity? | Hispanic or Latino(a); Not Hispanic or Latino(a); Unlisted ethnic minority; Prefer not to disclose |
| 8 | Select one | What is your age? | 25 or younger; 26-30; 31-35; 36 or older; Prefer not to disclose |
| 9 | Select one | Do you identify as any of the other following groups? [parent/caregiver, | Yes; No; Prefer not to disclose |

| | | | |
|---|---|---|---|
| | | non-US citizen or permanent resident, first in family to attend college, person with disability, veteran, neuro-atypical, started current program more than 5 years after completing most recent degree] | |
| 10 | Select one | If you identified as an underrepresented minority in any of the following aspects, do you feel that you and your advisor share any of those identities?<br>[woman or other gender minority, racial minority, sexual minority, parent/caregiver, non-US citizen or permanent resident, first in family to attend college, person with disability, veteran, neuro-atypical] | None of the above demographics; One of the above demographics; Two or more of the above demographics; Don't know |
| **II. EXPERIENCE WITH ADVISOR** | | | |
| **Question number** | **Question type** | **Text of question/ statement** | **Options provided to respondents** |
| 11 | Select one | How often do you meet with your advisor? | More than once a week; Weekly; 1-2 times per month; Less than once a month |
| 12 | Select one | How are meetings with your advisor scheduled? | Regularly scheduled; At my initiative; At my advisor's initiative; I can drop in without advance notice |
| 13 | Likert scale | My advisor is knowledgeable about my field and offers helpful advice on study design, research practices, data analysis, writing, and/or presentations. | 1 (Strongly Disagree), 2 (Disagree), 3 (Neutral), 4 (Agree), 5 (Strongly Agree), Don't Know, Not Applicable |
| 14 | Likert scale | My advisor is knowledgeable about preparation for academic careers. | 1 (Strongly Disagree), 2 (Disagree), 3 (Neutral), 4 (Agree), 5 (Strongly Agree), Don't Know, Not Applicable |
| 15 | Likert scale | My advisor is knowledgeable about preparation for non-academic careers. | 1 (Strongly Disagree), 2 (Disagree), 3 (Neutral), 4 (Agree), 5 (Strongly Agree), Don't Know, Not Applicable |

| | | | |
|---|---|---|---|
| 16 | Likert scale | I am satisfied with the frequency of meetings with my advisor. | 1 (Strongly Disagree), 2 (Disagree), 3 (Neutral), 4 (Agree), 5 (Strongly Agree), Don’t Know, Not Applicable |
| 17 | Likert scale | My advisor’s expectations of me are clear. | 1 (Strongly Disagree), 2 (Disagree), 3 (Neutral), 4 (Agree), 5 (Strongly Agree), Don’t Know, Not Applicable |
| 18 | Likert scale | My meetings are constructive and helpful in setting and achieving my goals. | 1 (Strongly Disagree), 2 (Disagree), 3 (Neutral), 4 (Agree), 5 (Strongly Agree), Don’t Know, Not Applicable |
| 19 | Likert scale | My advisor would advocate for me if needed. | 1 (Strongly Disagree), 2 (Disagree), 3 (Neutral), 4 (Agree), 5 (Strongly Agree), Don’t Know, Not Applicable |
| 20 | Likert scale | My advisor is supportive of my goals and ambitions. | 1 (Strongly Disagree), 2 (Disagree), 3 (Neutral), 4 (Agree), 5 (Strongly Agree), Don’t Know, Not Applicable |
| 21 | Likert scale | My advisor is empathetic to my concerns and issues. | 1 (Strongly Disagree), 2 (Disagree), 3 (Neutral), 4 (Agree), 5 (Strongly Agree), Don’t Know, Not Applicable |
| 22 | Likert scale | My advisor provides members of my lab with equitable access to opportunities for funding and/or collaboration. | 1 (Strongly Disagree), 2 (Disagree), 3 (Neutral), 4 (Agree), 5 (Strongly Agree), Don’t Know, Not Applicable |
| 23 | Likert scale | My advisor is transparent with members of my lab regarding decisions about funding, collaboration, and/or authorship. | 1 (Strongly Disagree), 2 (Disagree), 3 (Neutral), 4 (Agree), 5 (Strongly Agree), Don’t Know, Not Applicable |
| 24 | Likert scale | My advisor handles criticism and conflict well, and I would be comfortable bringing up issues that I have with them. | 1 (Strongly Disagree), 2 (Disagree), 3 (Neutral), 4 (Agree), 5 (Strongly Agree), Don’t Know, Not Applicable |
| 25 | Likert scale | My advisor handles conflict among lab members effectively, and I would be comfortable bringing issues to them. | 1 (Strongly Disagree), 2 (Disagree), 3 (Neutral), 4 (Agree), 5 (Strongly Agree), Don’t Know, Not Applicable |
| 26 | Likert scale | I am overall satisfied with the advising that I receive. | 1 (Strongly Disagree), 2 (Disagree), 3 (Neutral), 4 (Agree), 5 (Strongly Agree), Don’t Know, Not Applicable |
| 27 | Select | Are you co-advised?* | Yes; No |

| | | | |
|---|---|---|---|
| | one | *Students who responded "Yes" were directed to evaluate questions 12-26 for their secondary advisor. | |
| **III. RESEARCH GROUP** | | | |
| **Question number** | **Question type** | **Text of question/ statement** | **Options provided to respondents** |
| 28 | Select one | Other than yourself, how many graduate students, postdoctoral associates, or full-time research staff are associated with your lab? | Zero (only myself); 1-2; 3-5; 6-10; More than 10; Not sure |
| 29 | Select one | Including positions for research credit, work-study appointments, and volunteers, how many undergraduate students or part-time research staff are currently associated with your lab? | Zero; 1-5; 6-10; More than 10; Not sure |
| 30 | Select one | Are your thesis lab(s) affiliated with any of the following institutions? | Research museum (e.g., Museum of Vertebrate Zoology, University of California Museum of Paleontology, University and Jepson Herbaria, Essig Museum of Entomology); Research institute outside of your home department (e.g., Joint BioEnergy Institute, Plant Gene Expression Center, Joint Genome Institute, Lawrence Berkeley National Laboratory, Energy and Climate Institute); Graduate Group or Designated Emphasis outside of your home department (e.g. Graduate Group in Microbiology, Center for Computational Biology); None of the above |
| 31 | Likert scale | Resources within my thesis lab(s) are shared and/or distributed equitably (e.g. supplies/equipment, space, funding, research opportunities, time with advisor, networking opportunities). | 1 (Strongly Disagree), 2 (Disagree), 3 (Neutral), 4 (Agree), 5 (Strongly Agree), Don't Know, Not Applicable |
| 32 | Likert scale | When conflicts arise over resource distribution in my thesis lab(s), they are handled appropriately. | 1 (Strongly Disagree), 2 (Disagree), 3 (Neutral), 4 (Agree), 5 (Strongly Agree), Don't Know, Not Applicable |

| | | | |
|---|---|---|---|
| 33 | Likert scale | My research group encourages collaboration between lab members. | 1 (Strongly Disagree), 2 (Disagree), 3 (Neutral), 4 (Agree), 5 (Strongly Agree), Don't Know, Not Applicable |
| **IV. COMMUNITY AND CULTURE** | | | |
| **Question number** | **Question type** | **Text of question/ statement** | **Options provided to respondents** |
| 34 | Free response | Have you collaborated with any UC Berkeley faculty members in your home department other than your thesis advisor(s)? | N/A |
| 35 | Select one | Have you collaborated with any UC Berkeley faculty members outside of your home department? | More than once; Once; Never |
| 36 | Select one | Have you collaborated with postdoctoral associates, graduate students, or staff at UC Berkeley have you formally collaborated with on a research project? | More than once; Once; Never |
| 37 | Likert scale | I have received advice from UC Berkeley faculty members outside of formal instruction (e.g., courses, qualifying exam, committee) on research practices. | 1 (Strongly Disagree), 2 (Disagree), 3 (Neutral), 4 (Agree), 5 (Strongly Agree), Don't Know, Not Applicable |
| 38 | Likert scale | I have received advice from UC Berkeley faculty members outside of formal instruction on scientific careers. | 1 (Strongly Disagree), 2 (Disagree), 3 (Neutral), 4 (Agree), 5 (Strongly Agree), Don't Know, Not Applicable |
| 39 | Likert scale | I have received advice from UC Berkeley faculty members outside of formal instruction on issues relating to discrimination, physical or mental health, or other personal matters. | 1 (Strongly Disagree), 2 (Disagree), 3 (Neutral), 4 (Agree), 5 (Strongly Agree), Don't Know, Not Applicable |
| 40 | Likert scale | I have received advice from UC Berkeley graduate students, postdoctoral associates, or staff on research practices. | 1 (Strongly Disagree), 2 (Disagree), 3 (Neutral), 4 (Agree), 5 (Strongly Agree), Don't Know, Not Applicable |

| | | | |
|---|---|---|---|
| 41 | Likert scale | I have received advice from UC Berkeley graduate students, postdoctoral associates, or staff on scientific careers. | 1 (Strongly Disagree), 2 (Disagree), 3 (Neutral), 4 (Agree), 5 (Strongly Agree), Don't Know, Not Applicable |
| 42 | Likert scale | I have received advice from UC Berkeley graduate students, postdoctoral associates, or staff on issues relating to discrimination, physical or mental health, or personal matters. | 1 (Strongly Disagree), 2 (Disagree), 3 (Neutral), 4 (Agree), 5 (Strongly Agree), Don't Know, Not Applicable |
| 43 | Likert scale | I feel valued, included, and supported by members of my lab. | 1 (Strongly Disagree), 2 (Disagree), 3 (Neutral), 4 (Agree), 5 (Strongly Agree), Don't Know, Not Applicable |
| 44 | Likert scale | I feel valued, included, and supported by students in my graduate program. | 1 (Strongly Disagree), 2 (Disagree), 3 (Neutral), 4 (Agree), 5 (Strongly Agree), Don't Know, Not Applicable |
| 45 | Likert scale | If my lab is affiliated with a research museum, research institute, or graduate group outside of my home department, I feel valued, included, and supported by the members of the institute. | 1 (Strongly Disagree), 2 (Disagree), 3 (Neutral), 4 (Agree), 5 (Strongly Agree), Don't Know, Not Applicable |
| 46 | Likert scale | I feel valued, included, and supported by faculty and staff in my department. | 1 (Strongly Disagree), 2 (Disagree), 3 (Neutral), 4 (Agree), 5 (Strongly Agree), Don't Know, Not Applicable |
| 47 | Likert scale | My department fosters an environment that encourages collaboration between labs. | 1 (Strongly Disagree), 2 (Disagree), 3 (Neutral), 4 (Agree), 5 (Strongly Agree), Don't Know, Not Applicable |
| **V. STUDENT OUTCOMES** | | | |
| **Question number** | **Question type** | **Text of question/ statement** | **Options provided to respondents** |
| 48 | Select one | Have you presented a poster or talk at a formal annual society and/or national meeting? | More than once; Once; Never |
| 49 | Select one | Have you submitted a manuscript to a peer-reviewed journal? | More than once; Once; Never |

| | | | |
|---|---|---|---|
| 50 | Select one | Have you had a manuscript accepted or published in a peer-reviewed journal? | More than once; Once; Never |
| 51 | Likert scale | I am on track to complete my degree within normative time. | 1 (Strongly Disagree), 2 (Disagree), 3 (Neutral), 4 (Agree), 5 (Strongly Agree), Don't Know, Not Applicable |
| 52 | Likert scale | My research is meaningful and challenges me intellectually. | 1 (Strongly Disagree), 2 (Disagree), 3 (Neutral), 4 (Agree), 5 (Strongly Agree), Don't Know, Not Applicable |
| 53 | Likert scale | Travel or lab density restrictions imposed by COVID-19 reduced my research productivity during the past two years. | 1 (Strongly Disagree), 2 (Disagree), 3 (Neutral), 4 (Agree), 5 (Strongly Agree), Don't Know, Not Applicable |
| 54 | Likert scale | Personal or financial challenges imposed by COVID-19 reduced my research productivity during the past two years. | 1 (Strongly Disagree), 2 (Disagree), 3 (Neutral), 4 (Agree), 5 (Strongly Agree), Don't Know, Not Applicable |
| 55 | Likert scale | My program has adequately prepared me to pursue an academic research career. | 1 (Strongly Disagree), 2 (Disagree), 3 (Neutral), 4 (Agree), 5 (Strongly Agree), Don't Know, Not Applicable |
| 56 | Likert scale | My program has adequately prepared me to pursue a scientific career outside of academia. | 1 (Strongly Disagree), 2 (Disagree), 3 (Neutral), 4 (Agree), 5 (Strongly Agree), Don't Know, Not Applicable |
| 57 | Likert scale | I am happy and well-adjusted in my program. | 1 (Strongly Disagree), 2 (Disagree), 3 (Neutral), 4 (Agree), 5 (Strongly Agree), Don't Know, Not Applicable |
| 58 | Likert scale | I feel that I belong in my program. | 1 (Strongly Disagree), 2 (Disagree), 3 (Neutral), 4 (Agree), 5 (Strongly Agree), Don't Know, Not Applicable |
| 59 | Likert scale | Relative to other graduate students in my program, I would rate my progress on publications as… | 1 (Below Average), 2 (Average), 3 (Above Average), Don't Know |
| 60 | Likert scale | Relative to other graduate students in my program, I would rate my skill level at designing a research study as… | 1 (Below Average), 2 (Average), 3 (Above Average), Don't Know |
| 61 | Likert scale | Relative to other graduate students in my program, I would rate my skill | 1 (Below Average), 2 (Average), 3 (Above Average), Don't Know |

| | | | |
|---|---|---|---|
| | | level at applying for research funding as… | |
| 62 | Likert scale | Relative to other graduate students in my program, I would rate my skill level at executing a research study (data collection, model development, field survey) as… | 1 (Below Average), 2 (Average), 3 (Above Average), Don't Know |
| 63 | Likert scale | Relative to other graduate students in my program, I would rate my skill level at using software or programming languages typically needed in my field as… | 1 (Below Average), 2 (Average), 3 (Above Average), Don't Know |
| 64 | Likert scale | Relative to other graduate students in my program, I would rate my skill level at academic presentations as… | 1 (Below Average), 2 (Average), 3 (Above Average), Don't Know |
| 65 | Likert scale | Relative to other graduate students in my program, I would rate my skill level at communicating scientific results to the public as… | 1 (Below Average), 2 (Average), 3 (Above Average), Don't Know |
| 66 | Likert scale | Relative to other graduate students in my program, I would rate my skill level at leading and managing projects as… | 1 (Below Average), 2 (Average), 3 (Above Average), Don't Know |
| 67 | Likert scale | Relative to other graduate students in my program, I would rate my skill level at designing an academic course as… | 1 (Below Average), 2 (Average), 3 (Above Average), Don't Know |
| 68 | Likert scale | Relative to other graduate students in my program, I would rate my skill level at teaching a lab or discussion section as… | 1 (Below Average), 2 (Average), 3 (Above Average), Don't Know |
| 69 | Likert scale | Relative to other graduate students in my field, I would rate my progress on publications as… | 1 (Below Average), 2 (Average), 3 (Above Average), Don't Know |
| 70 | Likert scale | Relative to other graduate students in my field, I would rate my skill level at applying for research funding as… | 1 (Below Average), 2 (Average), 3 (Above Average), Don't Know |

| | | | |
|---|---|---|---|
| 71 | Likert scale | Relative to other graduate students in my field, I would rate my skill level at executing a research study (data collection, model development, field survey) as… | 1 (Below Average), 2 (Average), 3 (Above Average), Don't Know |
| 72 | Likert scale | Relative to other graduate students in my field, I would rate my skill level at using software or fieldming languages typically needed in my field as… | 1 (Below Average), 2 (Average), 3 (Above Average), Don't Know |
| 73 | Likert scale | Relative to other graduate students in my field, I would rate my skill level at academic presentations as… | 1 (Below Average), 2 (Average), 3 (Above Average), Don't Know |
| 74 | Likert scale | Relative to other graduate students in my field, I would rate my skill level at communicating scientific results to the public as… | 1 (Below Average), 2 (Average), 3 (Above Average), Don't Know |
| 75 | Likert scale | Relative to other graduate students in my field, I would rate my skill level at leading and managing projects as… | 1 (Below Average), 2 (Average), 3 (Above Average), Don't Know |
| 76 | Likert scale | Relative to other graduate students in my field, I would rate my skill level at designing an academic course as… | 1 (Below Average), 2 (Average), 3 (Above Average), Don't Know |
| 77 | Likert scale | Relative to other graduate students in my field, I would rate my skill level at teaching a lab or discussion section as… | 1 (Below Average), 2 (Average), 3 (Above Average), Don't Know |
